# Enabling Epitope Analysis of Glycoproteins via HDX-MS Using Antigens with Uniformed N-linked Glycans

**DOI:** 10.1101/2024.09.24.613422

**Authors:** Takeshi Ota, Masahiro Takayama, Takeshi Ishihara, Masaaki Sato, Sawaka Ono, Ryota Futamata, Masaya Fujitani, Masatomo Rokushima

## Abstract

Hydrogen deuterium exchange mass spectrometry (HDX-MS) is widely used for epitope analysis of the antibodies. However, epitope analysis of glycoproteins is challenging because of the heterogeneity of attached N-linked glycans. Recent studies have reported methods where N-linked glycans were removed at low pH following hydrogen deuterium exchange reactions, but the methods for glycoproteins remain controversial. Here, we demonstrate the utility of using antigens with uniformed N-linked glycans in glycoprotein epitope analysis by HDX-MS. By treating HEK293 cells with kifunensine, we were able to prepare antigens mainly with N-linked high-mannose-type glycans. Analysis of epitopes of a monoclonal antibody S309 using antigens prepared with this method allowed us to identify an epitope that included the previously reported N-linked glycan attachment sites for this antibody. We propose that using antigens with uniformed glycans should be effective for epitope analysis of glycoproteins, such as virus spike proteins covered by glycan shields. Moreover, we believe that this approach accelerates research on vaccines and neutralizing antibodies against viruses that escape host immunity through glycan shields or mutations.

## Introduction

Hydrogen-deuterium exchange coupled with mass spectrometry (HDX-MS) is commonly used for epitope mapping of antibodies. Recent examples include the use of HDX-MS to map the binding sites of monoclonal antibodies (mAbs) in a series of studies that led to the discovery of LY-CoV555^1^ and development of the REGEN-COV mAb cocktail,^2^ supplemented with cryo-electron microscopy structural analyses.

In a standard HDX-MS experiment, the target protein is incubated in quasi-neutral D2O buffer over a specified duration for replacement of hydrogen with deuterium. The sample was then diluted with an acidic quenching buffer without D2O. By maintaining a temperature close to 0°C and lowering the pH to approximately 2.7, the back-exchange of deuterated amides can be decelerated, thereby preserving deuteration. Thereafter, to achieve spatial resolution, the protein was digested using a protease at low pH to inhibit back-exchange reactions, and the peptides were analyzed by liquid chromatography and mass spectrometry (LC/MS). The rate of deuterium exchange at the peptide level was determined by quantifying the shift in the centroid *m/z* from the isotopic pattern of the undeuterated control peptide to the centroid *m/z* of the deuterated peptide using mass spectrometry.^3, 4^

In folded proteins, the deuterium uptake time of peptide backbone amides ranges from milliseconds to hours at physiological pH. This variation depends on the extent of involvement of the backbone amides in hydrogen bonding (e.g., in the preservation of secondary structures in proteins) or their isolation from the solvent. Hence, owing to protein-protein interactions, the rate of deuterium exchange changes not only at antibody-binding sites on the molecular surface, but also at allosteric sites. When implementing HDX-MS for antibody epitope analysis, the antigen-antibody binding interface restricts solvent accessibility, resulting in a reduced rate of deuterium exchange. This information allowed the identification of the epitope.

However, epitope analysis of glycoproteins using HDX-MS is challenging.^3^ This is because the high complexity of glycans leads to a reduction in the precursor signals of the peptides with glycans and their insufficient fragmentation. This makes glycopeptide identification difficult, and can cause problems with sequence coverage. Moreover, glycoproteomics, which is often used for glycopeptide identification, requires long separation times in liquid chromatography-tandem mass spectrometry (LC/MS/MS). In contrast, HDX-MS requires a shorter analysis time, thereby increasing ion suppression of peptides and making analysis even more difficult. In deuterium exchange reactions, the amide groups in the glycan structure can contribute to deuterium exchange, potentially complicating the measurement of deuterium uptake.^5^ These issues with glycoproteins are especially pronounced for viral spike proteins, which have evolved to evade human immune responses through acquisition of glycan shields and multiple glycan addition sites. In many HDX-MS studies, including those on SARS-CoV-2 and influenza, the areas around the N-linked glycan attachment sites were often excluded from the analysis.^2, 6^

Receptor-binding domain (RBD) of SARS-CoV-2 carries two N-linked glycans at N331 and N343. Several neutralizing antibodies that target the RBD, such as S309, have been reported to recognize epitopes that include the N343 glycan.^7^ This epitope exhibits high sequence conservation among different strains and contributes to a broad spectrum of neutralizing antibodies. Because of the critical importance of analyzing epitopes that include glycosylation sites in evaluating the mechanisms of neutralizing antibodies and vaccines for SARS-CoV-2, methodologies of HDX-MS analysis of spike proteins of this viral species have been studies. Recently, methods for enzymatically removing N-linked glycans after deuterium exchange have been developed as promising means to circumvent issues related to glycan modifications.^8^ However, these methods require an increase in temperature during the enzymatic glycan removal stage, leading to the inevitable reverse exchange of deuterium, which is debatable. Furthermore, these methods require the integration of new tools for glycan removal into existing HDX-MS systems, which could compromise their convenience.

In this study, we aimed to develop a general HDX-MS-based method for analyzing the epitopes of antibodies that work against glycoproteins, with the goal of elucidating the mechanisms and performance of neutralizing antibodies reacting to various virus spike proteins. To overcome the heterogeneity of N-linked glycans, we focused on the effects of kifunensine.^9^ This inhibitor halts the process of M9 high-mannose glycosylation by inhibiting the activity of Golgi mannosidase I in the Golgi apparatus. Furthermore, kifunensine has been reported to have a minimal impact on the quality control mechanisms of glycoproteins associated with endoplasmic reticulum-associated degradation^10^ and can be used for the production of homogeneously glycosylated recombinant proteins.^11^ Besides kifunensine, methods involving treatment with swainsonine have been reported,^9^ which inhibits Golgi mannosidase II or its production by cells lacking N-acetylglucosaminyltransferase I (GnTI).^12^ However, in terms of avoiding the generation of intermediate products during glycosylation, kifunensine was considered to be the most appropriate. Therefore, after preparing the antigen with kifunensine treatment and confirming the presence of high-mannose type glycans on the antigen by mass spectrometry, epitope analysis was performed using the HDX-MS system. Thus, we successfully developed a method to prepare antigens with homogeneous glycans and perform antibody epitope analysis without modifying the existing HDX-MS systems. Our method allowed us to achieve complete sequence coverage of the spike protein RBD and detect several binding regions of angiotensin converting enzyme-2 (ACE2) and mAb S309. In particular, the capability of the method was demonstrated by successful detection of the N343 glycan site of the RBD as an epitope of S309. These findings suggest the utility of our method to elucidate the epitopes of neutralizing antibodies that react with spike proteins of various viral strains.

## Experimental Section

### Recombinant Protein Synthesis for HDX-MS

Plasmids of recombinant SARS-CoV-2 spike protein consisting of a RBD, as described in UniProt ACC No. P0DTC2 and GenBank ACC No. QHD43416.1, and C-terminal conjugated 6xHis tag were transiently transfected into Expi293F cells using the ExpiFectamine 293 Transfection Kit, following the manufacturer’s guidelines (Thermo Fisher Scientific). After nurturing the cells for 4-5 days in the Expi293™ Expression Medium with kifunensine at a concentration of 5 - 20 μM,^9^ the supernatants were gathered and filtered by a 0.22-µm filter device (Merck Millipore). The TALON® Superflow Metal Affinity Resin and HisTALON™ Buffer Set (Takara Bio) was then used for recombinant protein purification from these supernatants as per the manufacturer’s directions. The eluates were concentrated using an Amicon Ultra-2 Centrifugal Filter Unit 30kDa NWMCO (Merck Millipore), in which the elution buffer was replaced with PBS (GIBCO).

### Binding Site Analysis

An HDX-PAL system (Leap Technologies) was used to automate the initiation of deuterium labeling, duration of the labeling reaction, quenching reaction time, injection into the UPLC system, and digestion time.^13^ A deuterium buffer (pH 7.4) was prepared using 10 mM PBS in D2O. For ACE2 protein binding analysis, a 1:1 molar ratio of recombinant spike protein and ACE2 protein (Product No. AC2-H5257; ACROBiosystems) were mixed and incubated at 37°C for 30 min to form complexes. For epitope mapping, a 1:1 molar ratio of the recombinant spike protein and antibody (clone No. S309, Abcam) was added and incubated at 37°C for 30 min to form complexes. Each of unbound and complexed samples was diluted 10-fold in either PBS, pH 7.4 (for non-deuterated experiments) or Deuterium buffer (for deuterated experiments) and was subjected to deuteration at 10°C with labeling periods of 60, 120, 240 s. The deuterium-labeled samples were then quenched at 0°C for 3 min by adding an equal volume of ice-cold quenching buffer (4 mol/L guanidine hydrochloride, 0.2 mol/L glycine hydrochloride, 0.5 mol/L TCEP, pH 2.7). The quenched samples were injected into an UltiMate 3000 UPLC system (Thermo Fisher Scientific). Online digestion was performed on an enzyme pepsin column (Waters) at 8°C for 270 s. The digested peptides were trapped on a Hypersil Gold column (Thermo Fisher Scientific) at 1°C and were then separated on an Acclaim PepMap300 C18 analytical column (Thermo Fisher Scientific) with a 7-minute gradient of 10%-35% B (mobile phase A: 0.1% formic acid in water, mobile phase B: 0.1% formic acid in acetonitrile). Mass spectra were acquired in the positive ion mode using an Orbitrap Eclips (Thermo Fisher Scientific) with parameters set at: electrospray voltage +4.0 kV, capillary temperature 275°C, full scan resolution 120000, MS/MS scan resolution 60000, *m/z* range 260-2000, scan time 1 s. For peptide identification, the recombinant proteins were analyzed under the aforementioned conditions, omitting the labeling reaction. All MS data were submitted to the ProteomeXchange Consortium via jPOSTrepo(https://repository.jpostdb.org/) using the dataset identifier, PXD056361/JPST003389.

### Peptide Identification

Software Byos 5.4.52 (Protein Metrics) was used for peptide identification with nonspecific cleavage, MS1 tolerance of 10 ppm, and MS2 fragment tolerance of 0.02 Da. Peptides and glycopeptides with a score of 40 or more were included in the final peptide set. Deamidation (at asparagine and glutamine) and oxidized methionine were used as dynamic modifications. N-linked glycans were dynamically modified using Byonic’s usual modification.^14^

### Data analysis of hydrogen/deuterium exchange

HDExaminer software, version 3.1.0 (Sierra Analytics) was used to determine the hydrogen/deuterium exchange ratio for each identified peptide based on the raw MS data files from all HDX experiments.^15^ To avoid the complexity of the data analysis, when peptides of the same sequence were identified at different charge states, their charge states were standardized to a single state using software. The parameters for the HDX-MS analysis are listed in Supplemental Table 1. Subsequently, all peptides analyzed using the HDExaminer were manually refined to decrease false-positive results. The deuteration differences (Δ%D) between unbound and complexed samples were computed for each peptide. Peptides that met criteria 1 and 2 were selected.^2^ When calculating the Δ%D of the 3-fold value of the Standard Deviation for the peptide length excluding the terminal 2 amino acids from the average detected peptide length, the value was approximately 1.5%. This indicates that Criterion 1 is sufficiently stringent.

Criterion 1: Δ%D is greater than 5% for two or more adjacent peptides.

Criterion 2: The peptide lengths are 4 amino acids or more.

### Visualization of Protein Binding Sites on the RBD Structure

The data stored in public protein structure databases often lack details regarding the structures of protein surfaces or loops. Hence, AlphaFold version 2.2.2 (DeepMind) was used to model the RBD structure.^16^ Modeling was conducted using protein information up to September 1, 2022, as a reference template. Additionally, the protein-binding site and its surrounding structures were clearly illustrated using PyMOL version 2.5.5 (Schrödinger).

## Results and Discussion

### Peptide Mapping of kifunensine-treated RBD

In reports on the peptide mapping of glycoproteins, glycopeptides are considered more difficult to identify than non-glycosylated peptides in LC/MS/MS. An analytical software (Byonic) that enables glycopeptides identification has been developed in recent years.^14^ Although we used this software, the identification efficiency for glycopeptides was lower than that for non-glycosylated peptides (data not shown). We speculate that this is mainly due to the heterogeneity of the glycans and the subsequent weakening of the signal in mass spectrometry. Therefore, we treated cells with kifunensine during protein expression to produce a recombinant RBD with homogeneous glycans. Subsequently, the sample was digested with pepsin, and peptide mapping was performed using LC/MS/MS in several measurement ranges and charge state combinations, which enabled the identification of peptides with a 100% amino acid sequence coverage rate (Figure 1A). Performing LC/MS/MS under several conditions had a consistent effect on improving peptide identification scores by Byos (data not shown). Regarding the glycan structure of the two glycosylation sites (N331 and N343), we individually examined if the high-mannose-type glycan M9 was the main structure. As expected, we could confirm that almost uniform glycosylation occurred through the inhibition of class I alpha-mannosidase by kifunensine. For the RBD prepared using kifunensine, the minimum amount of protein required for 100% amino acid sequence coverage was lower than that of the RBD prepared without kifunensine (data not shown). This suggests that converging glycans to the high-mannose type dramatically improves glycopeptide detection sensitivity.

**Figure 1.**
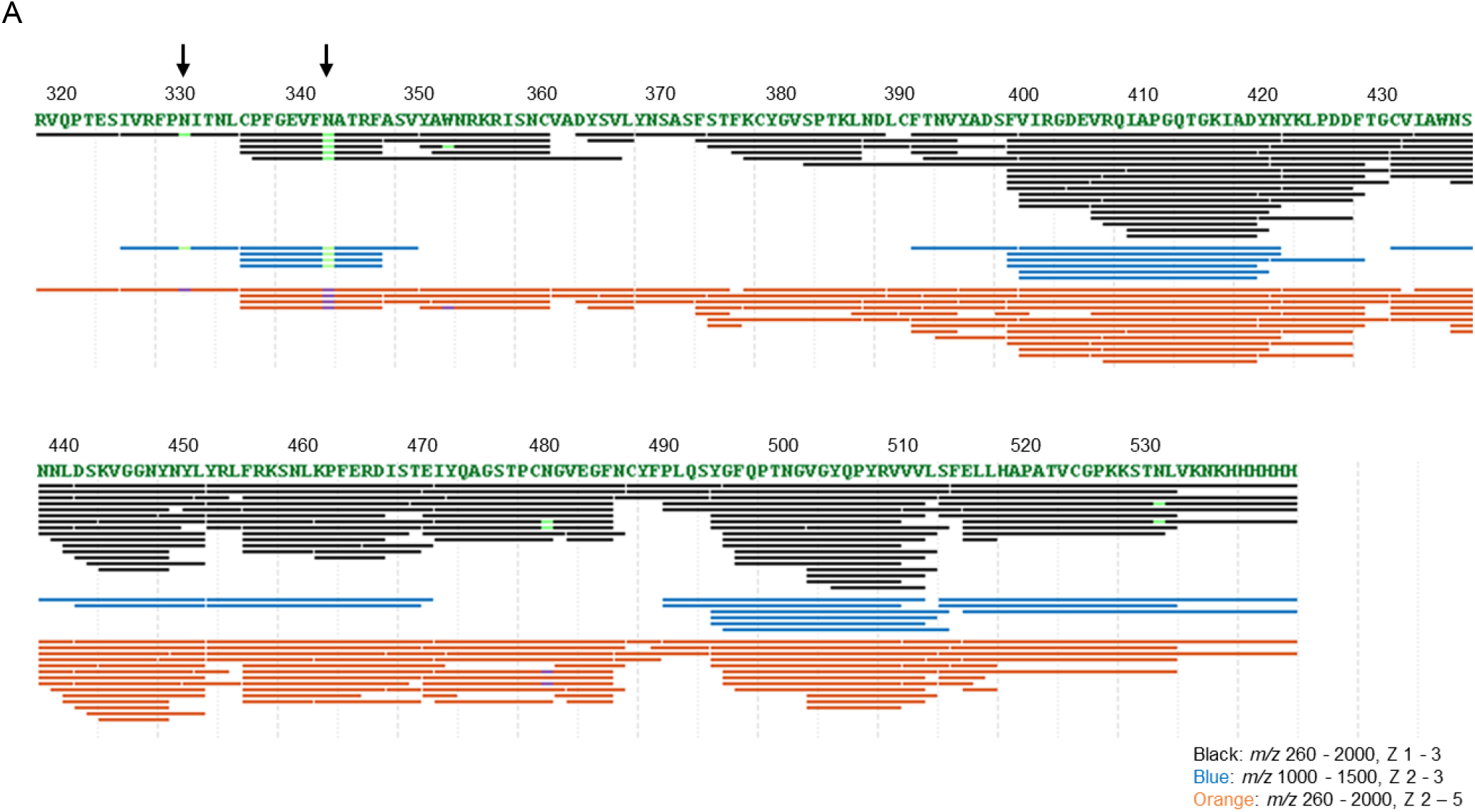
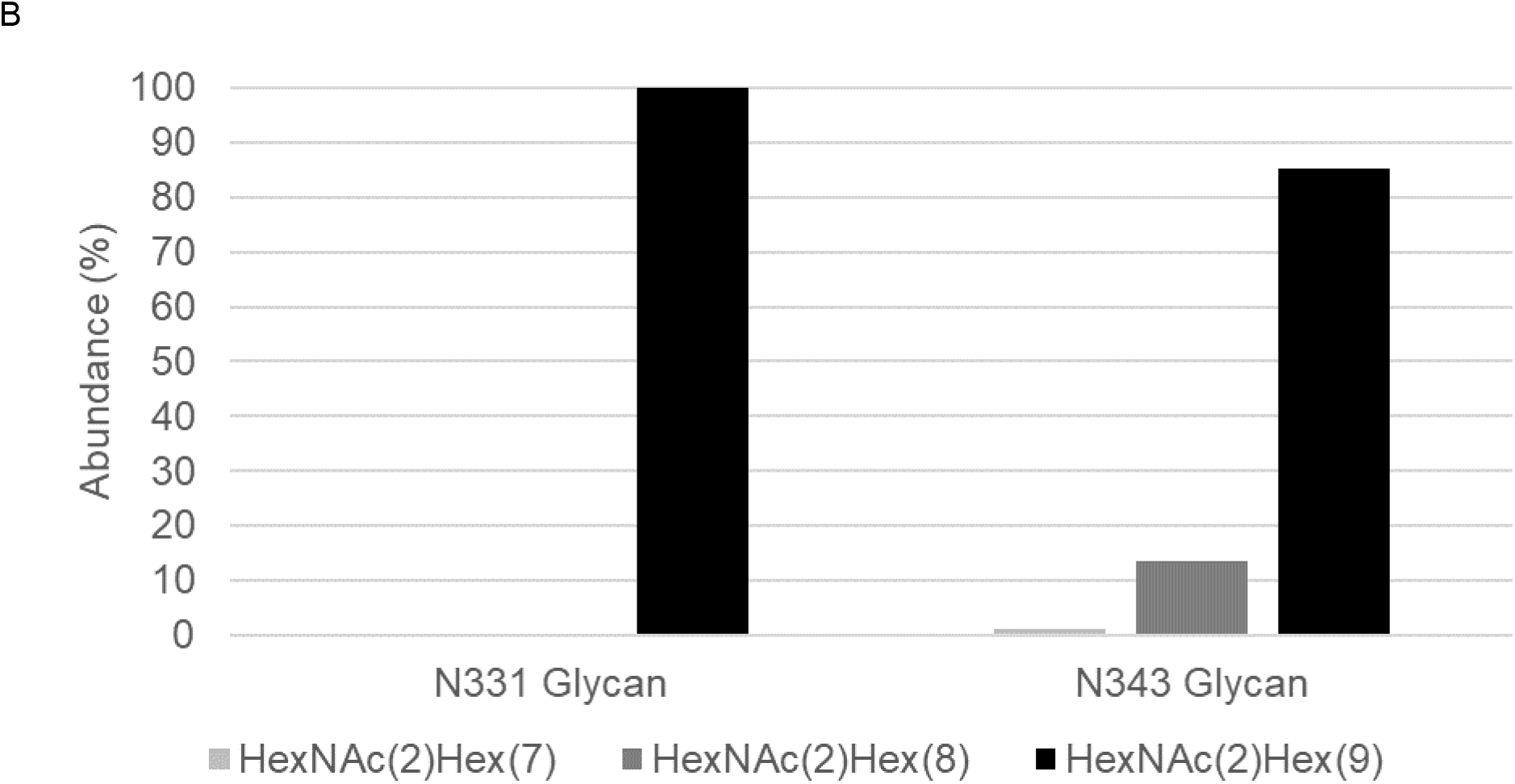
Peptide Mapping of kifunensine-treated RBD The kifunensine-treated RBD was digested with pepsin and analyzed by LC/MS/MS. (A) The sequence number of amino acids starts from the 319th of the wild-type full-length spike protein. The site of N-linked glycan addition is indicated by arrows. The length of the bar represents the length of the peptide. Black, blue, and orange represent different conditions of mass spectrometry measurement. Within the bar, the spots where the color is inverted (actually, black to yellow-green, blue to yellow-green, orange to blue) represent post-translational modifications (N-linked glycan modification or deamidation). (B) The proportion of the N-linked glycan structures added to N331 and N343 was calculated from the area values of the glycopeptides identified by LC/MS/MS. The sum of the values for each site equals 100%.

However, while the non-labeled N-terminal peptides of the position 319-336 could be easily identified through peptide mapping, it was often difficult to detect deuterium-labeled N-terminal peptides of that position in HDX experiments (Supplemental Figure 2). This may be due to the effects of the signal sequence artificially conjugated to RBD, or the poor compatibility between the amino acid sequence of the RBD and pepsin digestion.

### Analysis of the Binding Site of High-Mannose Type RBD and ACE2 by HDX

It has been reported that RBD binds to ACE2.^17^ Meanwhile, it is suggested that differences in glycan structure may alter protein conformation, potentially impacting its function.^18^ Therefore, we performed binding site analysis with ACE2 to investigate whether the recombinant RBD, to which we added high-mannose-type glycans (hereafter called high-mannose-type RBD), forms a functional conformation. After incubating the high-mannose RBD with recombinant ACE2 at a 1:1 molar ratio, we performed binding site analysis using HDX-MS and obtained results similar to those previously reported.^19^ Regions with decreased deuterium exchange rates were divided into four; 388-395, 432-453, 470-487, and 491-514 amino acids, and four or more peptides with reduced exchanges were detected in each region (Figure 2A). In addition, a time-dependent perturbation in the deuterium exchange rate was examined up to 240 s (Figure 2B). However, we did not detect any changes in the deuterium exchange rate for peptides including K417, L455, F456, Y489, and F490, which are among ACE2 binding sites reported previously (Supplemental Figure 1).^19^ We presume that the incorporation rate of deuterium in each peptide, including K417, L455, F456, Y489, and F490, was low owing to the influence of the secondary structure and conformation. Consequently, we could not detect the difference in deuterium incorporation rate between the presence and absence of ACE2.

**Figure 2.**
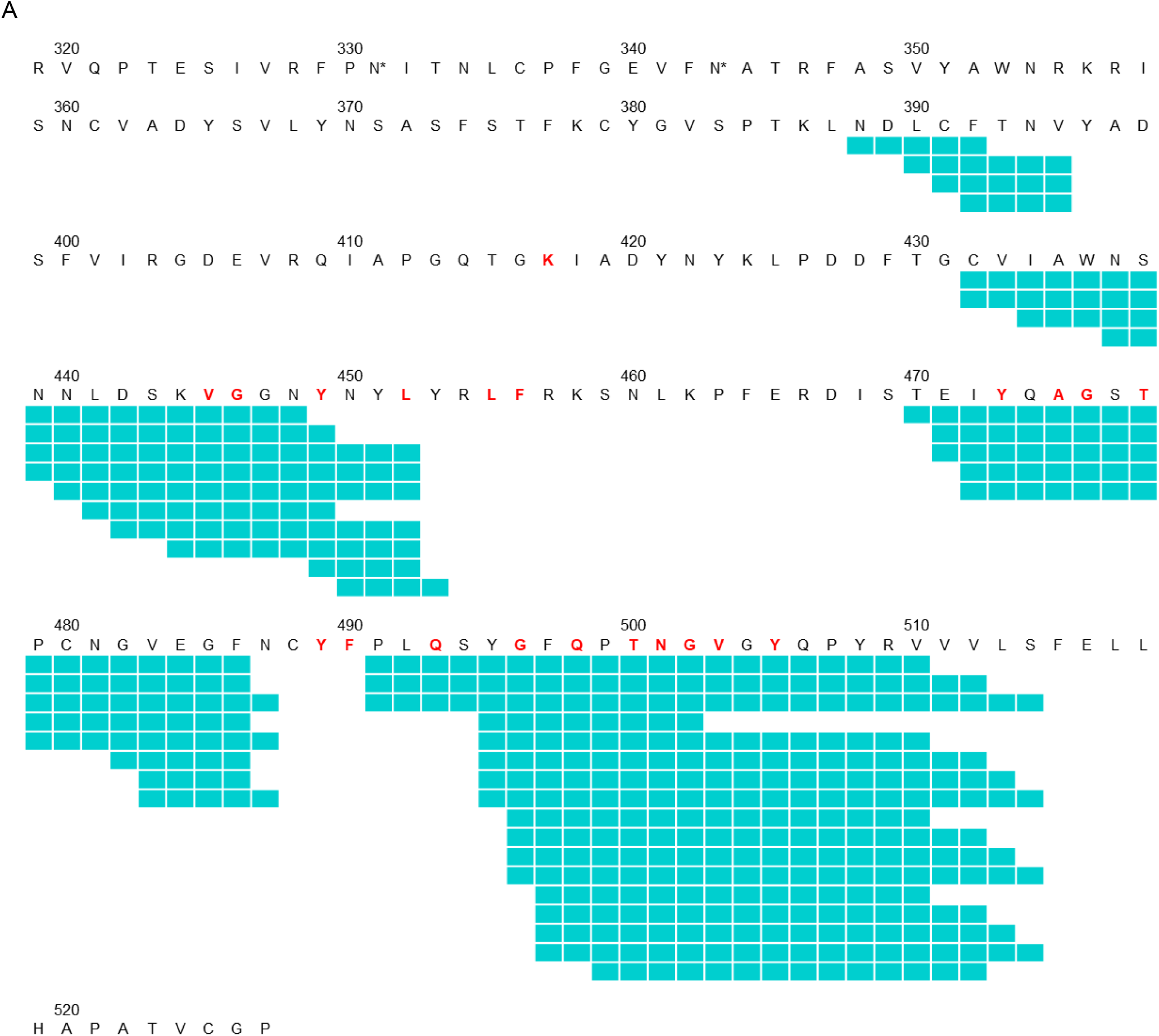
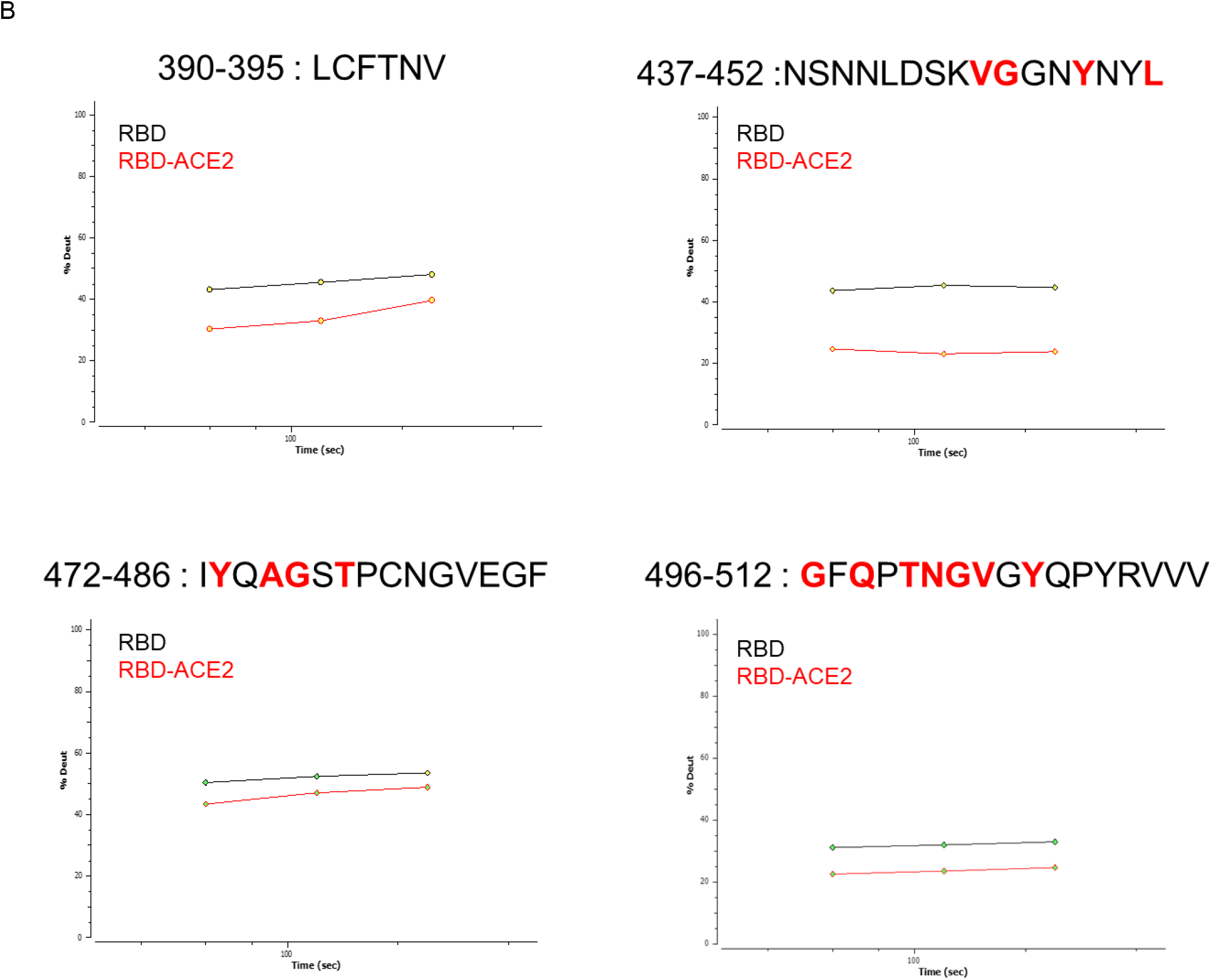
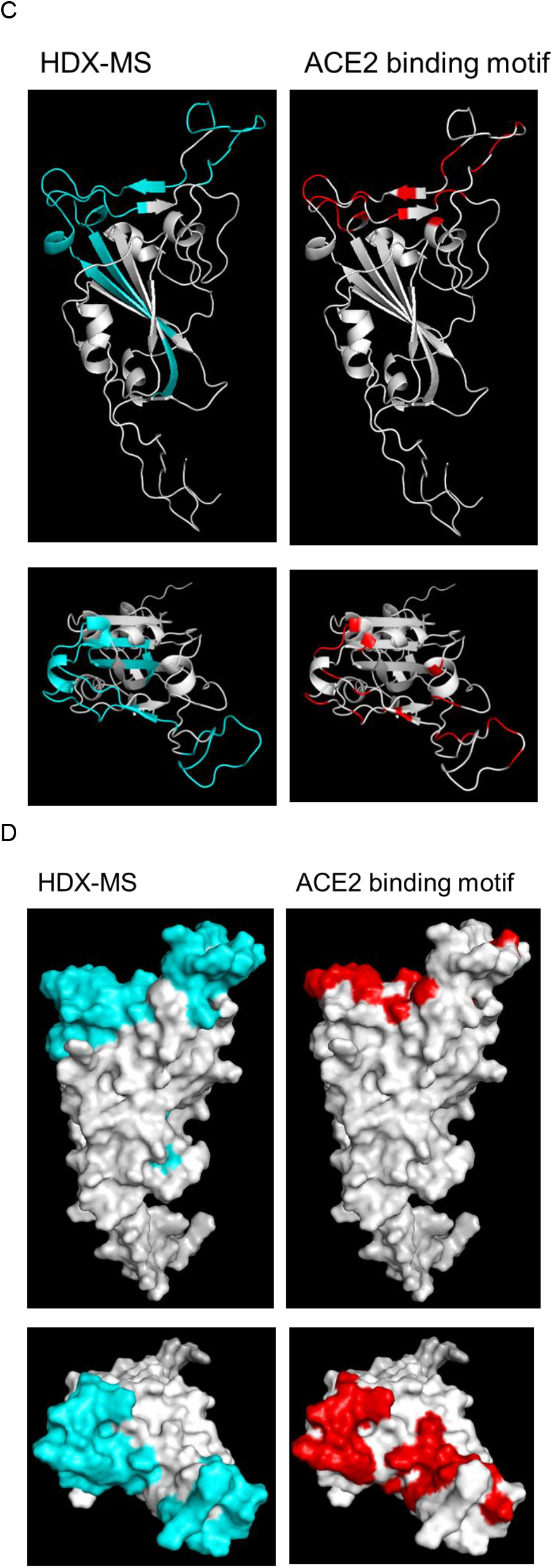
HDX analysis of ACE2 binding site on high-mannose type RBD The high-mannose-type RBD was analyzed using HDX-MS in the presence and absence of ACE2. (A) This sequence exhibits amino acids from the 319th to the 527th position in the wild-type full-length spike protein. Asterisks represent N-glycosylation sites. Red bold letters indicate the previously reported ACE2 binding amino acids. The light green bar, composed of consecutive rectangles, indicates the location of the peptides that fulfilled the criteria for reduced deuterium uptake. (B) Graphs show the temporal changes in the uptake of deuterium by representative peptides, where the incorporation of deuterium decreased upon binding with ACE2 in high-mannose type RBD. The black line represents the condition of RBD alone, whereas the red line represents the condition of RBD in the presence of ACE2. The x-axis denotes time and the y-axis denotes the amount of deuterium uptake. (C, D) Merged peptides obtained from figure 2A (light green) and previously reported ACE2 binding amino acids (red) were visualized in the structural model of the RBD. The lower panel represents a 90-degree forward rotation of the structure depicted in the upper panel. C is a ribbon diagram, and D is a surface diagram.

When regions with decreased deuterium exchange rates were shown in the three-dimensional structure, there were significantly different regions between the HDX-MS results and those of previous report on ACE2 biding motif (Figure 2C and 2D).^19^ By comparing the data of our study and the previous report, we found that regions 388-395, 432-440, 479-487 and 507-514 amino acids, where the deuterium exchange rate differed between them, were located inside the three-dimensional structure of the RBD. Considering the position of each region the three-dimensional structure, we assumed that the deuterium exchange rate in 388-395 amino acids decreased due to an allosteric conformational change in the RBD by ACE2 binding, as they are located on the opposite face from the ACE2 binding site. In addition, the regions of 432-440 and 507-514 amino acids, which are not among reported ACE2 binding sites, are included in the detected peptides with decreased deuterium exchanges (Figure 2A). Because these regions area located in the beta-sheet in the center of RBD and their exposure to the solvent is limited, they are considered not to be directly involved in ACE2 binding. This result should reflect relatively low-resolution in epitope analysis, which is due to the long peptides produced by pepsin digestion. Similarly, the region of 479-487 amino acids may not have had a direct role in ACE2 because of either restricted access to the solvent by binding of ACE2, or a conformational change (Figure 2A). However, although there were some discrepancies, when exclusively focused on the region of protein surface, the HDX-MS findings were in good agreement with those of previous reports (Figure 2D). Based on these observations, we concluded that the high-man-nose-type RBD forms a functional conformation.

### Analysis of the Binding Site of High-Mannose Type RBD and S309 by HDX

Figure 2 shows that the high-mannose type RBD forms a functional conformation. Therefore, we conducted a binding-site analysis of the monoclonal antibody S309, which uses the glycosylation site as a part of its epitope. Similar to the analysis of the ACE2 binding site, we incubated the high-mannose type RBD and S309 at a 1:1 molar ratio and conducted binding site analysis using HDX-MS. Our results were similar to those previously reported.^7^ The areas where the deuterium exchange rate decreased were divided into three regions, consisting of 336-347, 348-365, and 432-443 amino acids, and two or more peptides with varied length and/or positions were detected in each area (Figure. 3A). A constant decrease in the deuterium exchange rate was observed for up to 240 s for the hydrogen-deuterium exchange reaction in the presence of S309 (Figure. 3B). Additionally, a decrease in deuterium uptake was detected in the 388-395 amino acids located near the three regions described above (Supplemental Figure 2). This region has not been reported in cryoelectron microscopy observations and may be an epitope created by the fluctuation of S309 binding to the RBD. ^7^

**Figure 3.**
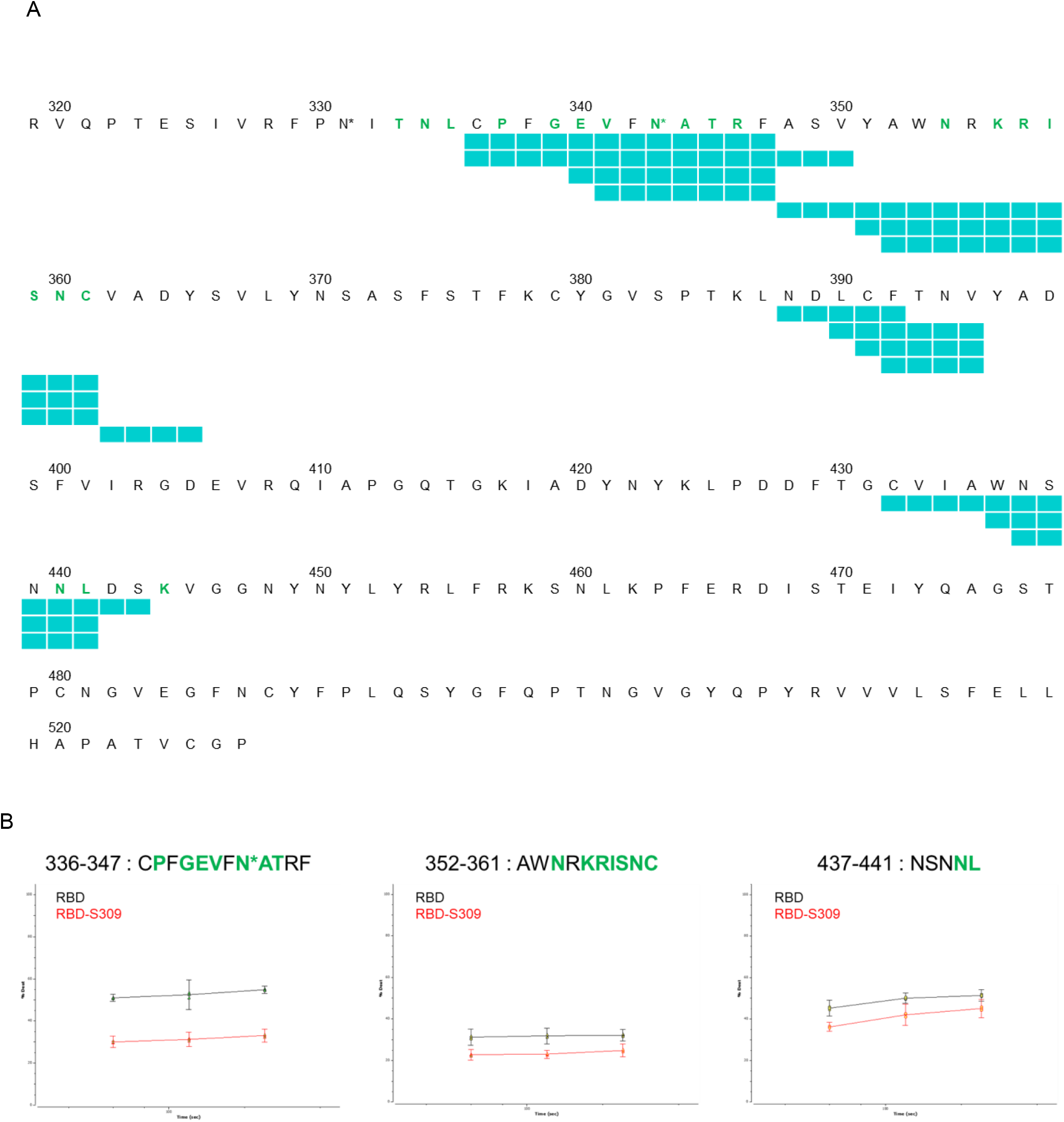
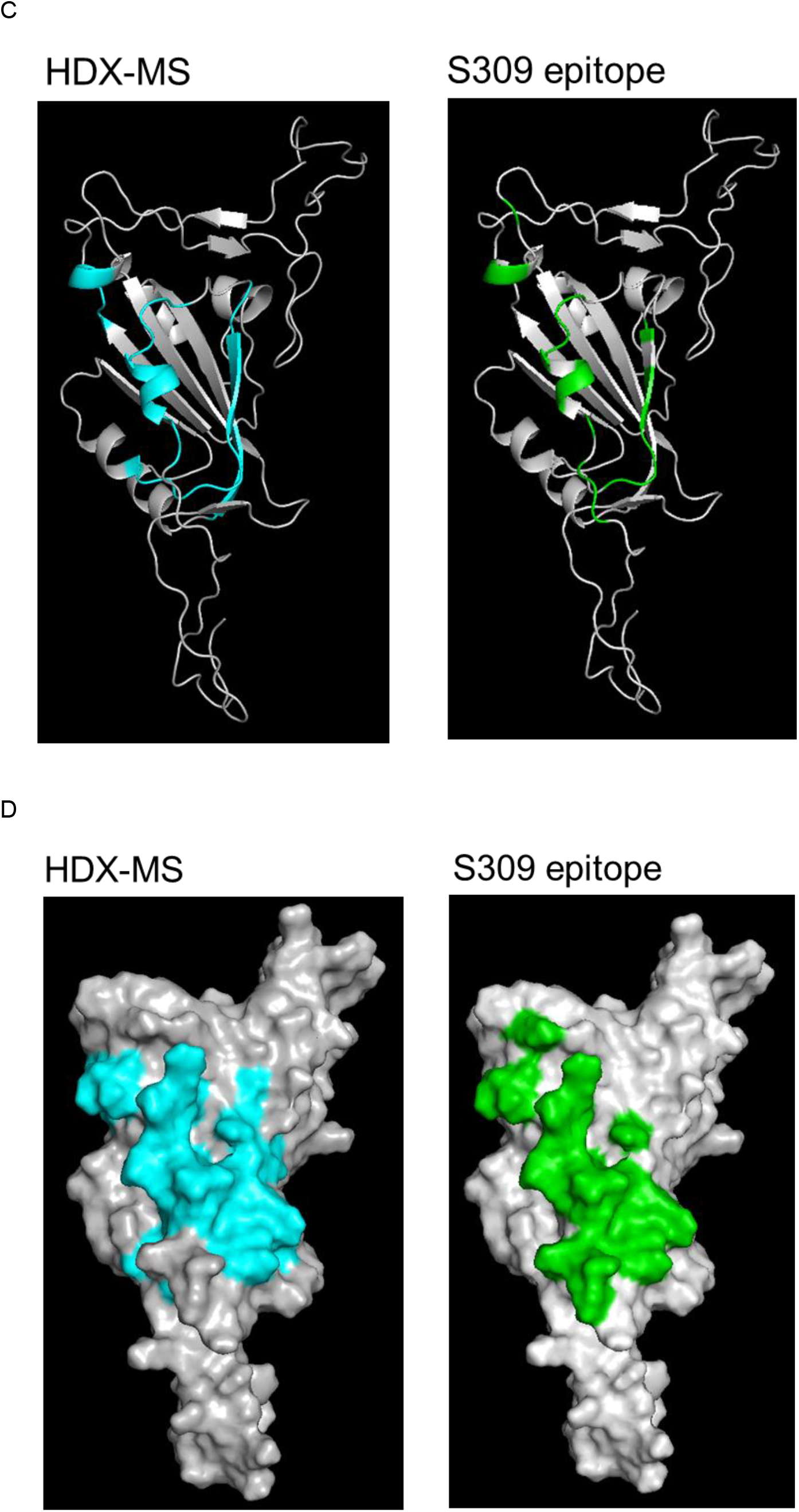
HDX analysis of S309 binding site on high-mannose type RBD High-mannose type RBD were analyzed by HDX-MS in the presence and absence of S309. (A) The sequence, asterisk, and light green bars have the same meaning as in figure 2A. Green bold letters indicate the previously reported S309 binding amino acids. (B) Graphs show the temporal changes in the uptake of deuterium by representative peptides, where the incorporation of deuterium decreased upon binding with S309 in high-mannose type RBD. The black line represents the condition of RBD alone, whereas the red line represents the condition of RBD in the presence of S309. The x-axis denotes time and the y-axis denotes the amount of deuterium uptake. (C, D) Merged peptides obtained from figure 3A (light green) and previously reported S309 binding amino acids (green) were visualized in the structural model of the RBD. C is a ribbon diagram, and D is a surface diagram.

Regarding other reported epitopes of S309, we could not identify peptides including the T333, N334, and L335 epitopes of S309. This may be due the signal peptide artificially attached to RBD and/or the poor compatibility between the RBD sequence and pepsin digestion (mentioned above). Furthermore, we did not detect any changes in the deuterium exchange rate of the peptides that included the K444 epitope of S309 with unknown reason (Supplementary Figure 2).

For the peptide (336-347 amino acids) that is glycosylated and included in the epitope of S309, the deuterium exchange rate was over 50% in the absence of S309, which was higher than that of the other peptides, probably because of the influence of deuterium exchange on glycosylation. Nonetheless, we detected a decrease in the deuterium exchange rate in the presence of S309. When the regions where the deuterium exchange rate decreased were illustrated in the three-dimensional structure, it was confirmed that the epitopes identified in this study were consistent with those of the previous reports (Figure 3C and 3D).^7^ From these results, we confirmed that it is possible to properly monitor the hydrogen-deuterium exchange reaction at glycosylation sites (Figure 3B) and demonstrated the utility of our method for epitope analysis of antibodies that recognize glycosylation sites.

## Conclusions

In this study, we prepared antigens in a mammalian cell protein expression system with the use of kifunensine, which inhibits the processing of N-linked glycans, thereby converting various glycans into high-mannose type glycans. We then demonstrated that the interaction sites of antibodies recognizing glycans and their vicinity could be identified as epitopes using these antigens. Therefore, we believe that we developed a potentially universal methodology for epitope analysis of antibodies binding to glycoproteins.

In previous studies, epitope analysis of antibodies binding to glycoproteins often excluded the areas around the glycosylation sites because of glycan issues inherent in HDX-MS.^6^ This is particularly true for spike proteins of viruses, such as SARS-CoV-2, which have evolved to evade human immune responses through glycan shields. One potential solution is the use of a protein expression system that does not involve glycan modification during sample preparation. However, care must be taken as glycans can significantly influence various processes. For example, there may be difficulties in protein expression itself when using systems such as *Escherichia coli* or cell-free glycoprotein synthesis systems. Furthermore, even when a glycoprotein is expressed, it is uncertain whether it adopts the correct conformation.^20^ Another solution may involve removing glycans from the glycoprotein prior to HDX-MS, but it is unclear whether glycans can be completely removed. Moreover, the effects of glycan removal on the conformation must also be considered. For instance, the spike protein of wild-type SARS-CoV2 has 22 N-linked glycan sites,^21^ and it is challenging to fully remove these glycans under native conditions, rendering this approach impractical.

Recently, the N-linked glycan-removing enzymes, PNGase Dj and Rc, which retain their enzymatic activity even under acidic conditions, have been discovered. As a result, a method has been developed in which the exchange reaction with deuterium is initiated for the glycoprotein, the reaction is stopped at a low pH, the glycoprotein is digested in a pepsin-immobilized column, followed by the removal of the glycans with a PNGase Dj-immobilized column, and then HDX-MS is performed.^8, 22^ However, this method remains controversial, as the temperature temporarily increases during the glycan removal process, which may lead to signal reduction due to back exchange. In this study, we enabled the epitope analysis of glycoproteins without removing glycans, in which all glycan attachment sites were converted into high-mannose-type glycans. Importantly, this method does not incorporate steps that might increase deuterium back-exchange, and alterations to the HDX-MS system are not required. This allows for the antigen-antibody epitope analysis of both glycoproteins and non-glycosylated proteins, which is considered a significant advantage of this method.

Our method does have several potential limitations. Firstly, the transformation of glycans to the high-mannose type could hinder pepsin digestion, although we could not observe any decrease in sequence coverage or noticeable impact on enzymatic digestion in this study (Figure 1A). Secondly, the acetamide group of glycans in glyco-peptides often shows high deuterium exchange rates, which can interfere with the measurement of deuterium uptake.^5^ Nevertheless, this effect did not impede the epitope analysis in this study (e.g., 336-347 amino acids in Figure 3B). Thirdly, high-mannose type glycans potentially influence intramolecular conformation and protein-protein interactions such as those between antibody and antigen. However, in our study, these effects seemed minimal because we confirmed maintained binding ability of RBD with ACE2 and obtained the results of epitope analysis for S309 similar to previous studies. On the other hand, as it has been reported that glycans do influence protein conformation, the antigen with uniformed glycans may have limited applicability in researches related to conformational dynamics.^18, 23^

Our method should offer a universal solution for epitope analysis of antibodies binding to glycoproteins. Notably, broad and potent neutralizing antibodies that recognize glycosylated epitopes have been reported in the spike proteins of viruses with glycan shields, such as SARS-CoV2^24^ and HIV.^25^ This makes antibody epitope analysis particularly valuable. Moreover, the spike proteins of currently prevalent viruses like SARS-CoV2 and influenza, both of which are glycoproteins, exhibit rapid mutation rates. In contrast, X-ray crystallography and cryo-electron microscopy analyses can elucidate an epitope at the atomic level but are often laborious because of low throughput.^3^ This does not necessarily lend itself well to the epitope analysis of antibodies against spike proteins (all glycoproteins) with various mutations, particularly in viruses such as SARS-CoV2 and influenza, which are currently prevalent and mutate rapidly.

To address some of these issues, we propose combining our method with protein structure prediction (e.g., using AlphaFold2 software to predict the structure of mutated spike proteins). This should rapidly facilitate epitope analysis of neutralizing antibodies derived from humans or mice, and consequently, enables prediction of the performance of these antibodies against mutated viruses. This method can contribute to the development of therapeutics such as vaccines and antibody drugs that are resistant to virus mutations.^26-28^

## Author Information

### Present Addresses

Shionogi Pharmaceutical Research Center, 3-1-1 Futaba-cho, Toyonaka, Osaka 561-0825, Japan.

## Author Contributions

Takeshi Ota and Masahiro Takayama designed the study. Masatomo Rokushima contributed to the development and validation of the system. Takeshi Ishihara, Masaaki Sato and Ryota Futamata designed and prepared of proteins. Masaya Fujitani prepared structural model of antigen. Masahiro Takayama and Sawaka Ono performed HDX-MS and data analysis. Takeshi Ota, Masahiro Takayama and Masatomo Rokushima wrote the manuscript.

## Notes

The authors declare no conflicts of interest associated with this manuscript.

## Acknowledgment

The authors thank all our colleagues who participated in the COVID-19 vaccine program at Shionogi. The authors thank Ueda, Hiroshi (Shionogi & Co., Ltd.) for their support in writing this paper. We would like to thank Editage (www.editage.jp) for English language editing.

## Associated content

Supporting Information

**Supplemental Table 1.** Data acquisition and analysis parameters for the HDX-MS experiment on the RBD and RBD-S309 complex.

| Data set | RBD<br>(control) | RBD +<br>S309 |
| --- | --- | --- |
| Back-exchange (mean / IQR over entire project) | unknown |  |
| # of peptides | 148 | 148 |
| Sequence coverage | 92.09% | 92.09% |
| Average peptide length / Redundancy | 12.38 / 8.52 | 12.38 / 8.52 |
| Replicates | 3 | 3 |
| Repeatability (avg. stddev of #D) | 0.0538 | 0.0528 |
| Significant differences in HDX (delta HDX > X D - 99% CI) | n/a | 0.4251 D |
| 3SD |  | 0.1584 |
| $\Delta\%D$ of 3SD ($(3SD/(\text{Average peptide length}-2)*100)$) | | 1.526% |

**Supplemental Figure 1.**
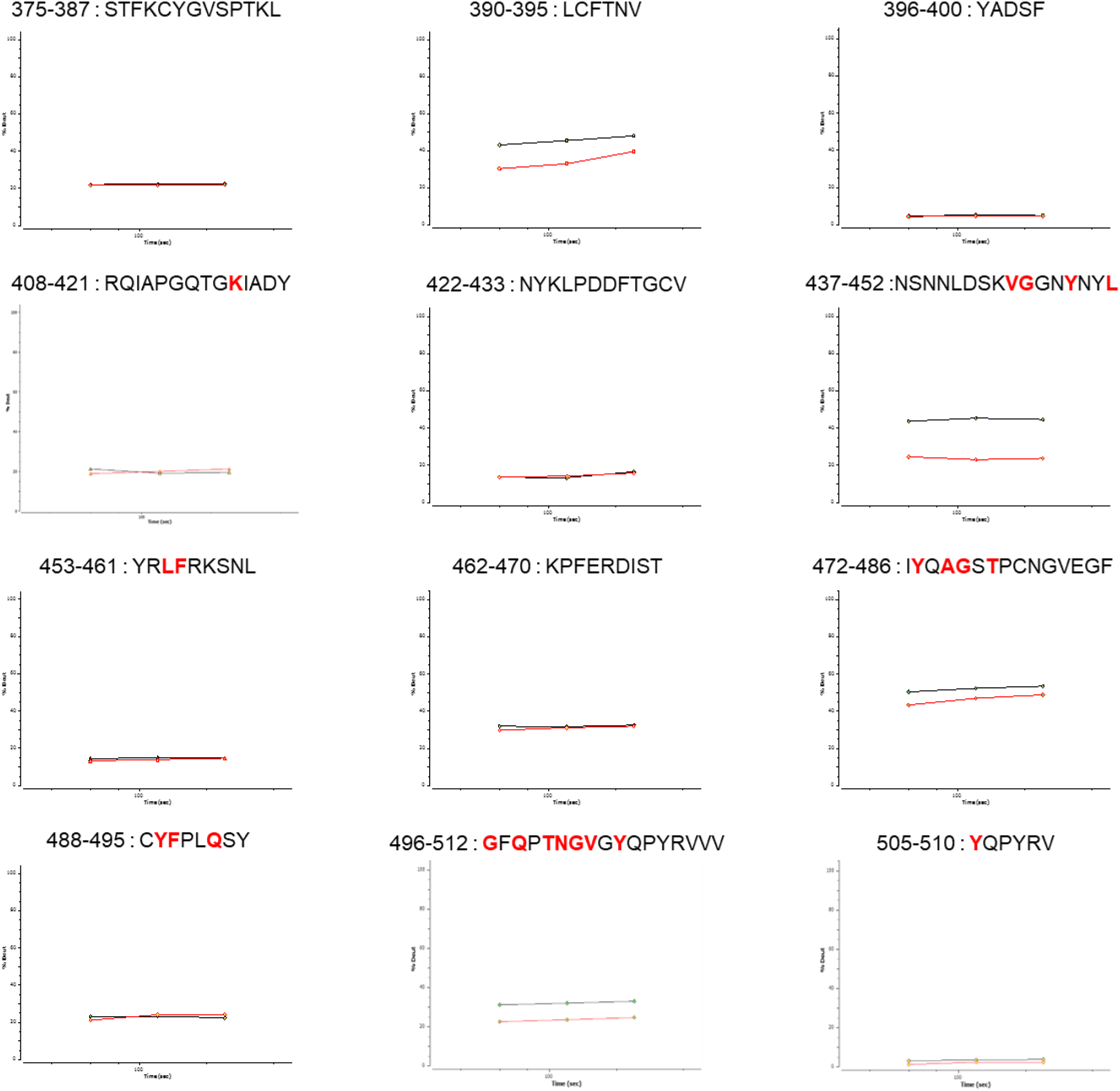
This figure presents data from Figure 2B, supplemented with HDX-MS data on adjacent peptides. The black line represents the condition of RBD alone, whereas the red line represents the condition of RBD in the presence of ACE2. The x-axis denotes time and the y-axis denotes the amount of deuterium uptake. Red bold letters indicate the previously reported ACE2 binding amino acids.

**Supplemental Figure 2.**
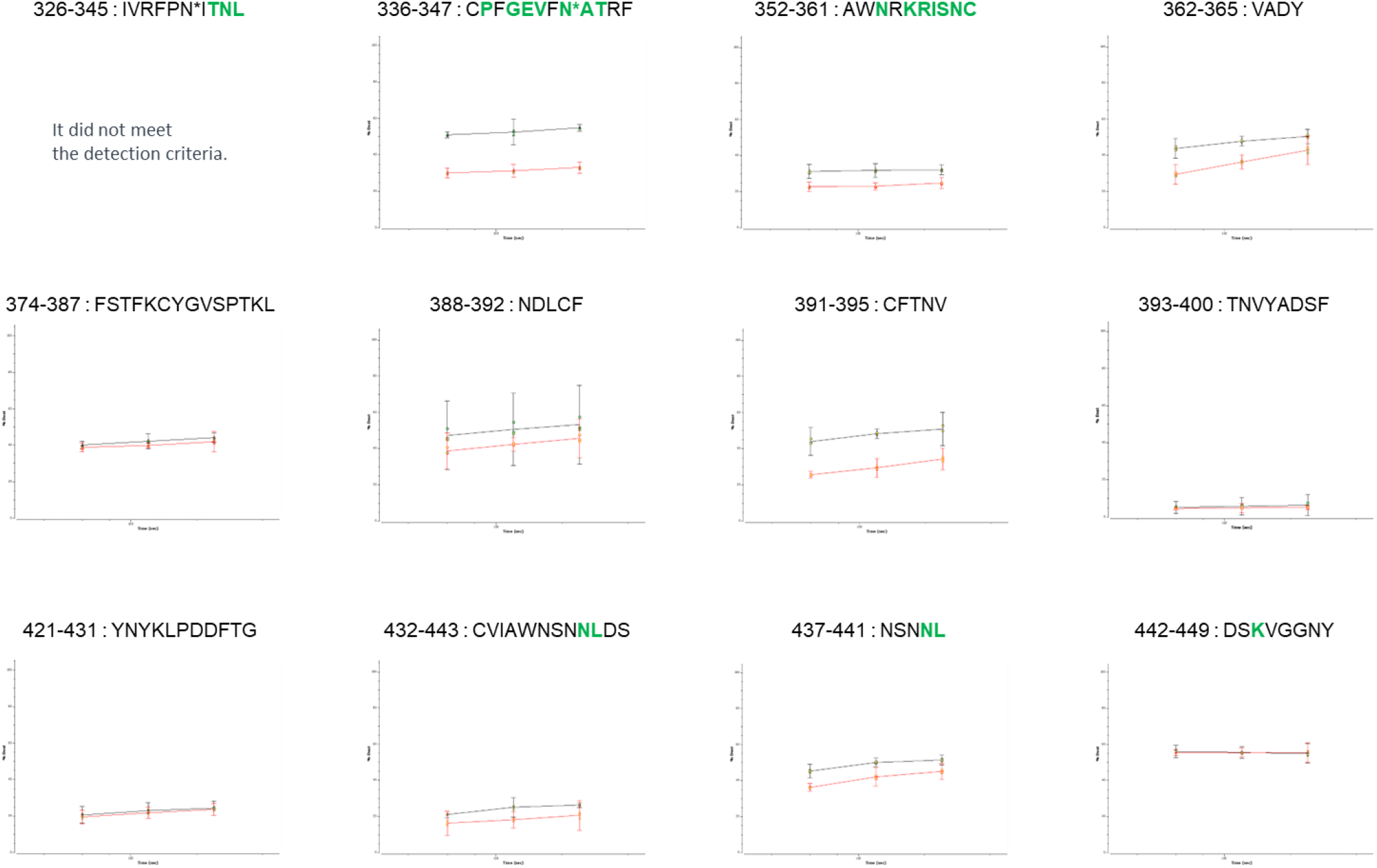
This figure presents the data from figure 3B supplemented with HDX-MS data on adjacent peptides. The black line represents the condition of RBD alone, whereas the red line represents the condition of RBD in the presence of S309. The x-axis denotes time and the y-axis denotes the amount of deuterium uptake. Green bold letters indicate the previously reported S309 binding amino acids.

